# Identification of an immune gene expression signature associated with favorable clinical features in Treg-enriched patient tumor samples

**DOI:** 10.1101/246603

**Authors:** Kevin B. Givechian, Kamil Wnuk, Chad Garner, Stephen Benz, Hermes Garban, Shahrooz Rabizadeh, Kayvan Niazi, Patrick Soon-Shiong

**Author notes:** **Address all correspondence to:** Kevin B. Givechian, NantOmics LLC, Culver City, CA 90232, USA.

## Abstract

Immune heterogeneity within the tumor microenvironment undoubtedly adds several layers of complexity to our understanding of drug sensitivity and patient prognosis across various cancer types. Within the tumor microenvironment, immunogenicity is a favorable clinical feature in part driven by the antitumor activity of CD8+ T cells. However, tumors often inhibit this antitumor activity by exploiting the suppressive function of Regulatory T cells (Tregs), thus suppressing the adaptive immune response. Despite the seemingly intuitive immunosuppressive biology of Tregs, prognostic studies have produced contradictory results regarding the relationship between Treg enrichment and survival. We therefore analyzed RNA-seq data of Treg-enriched tumor samples to derive a pan-cancer gene signature able to help reconcile the inconsistent results of Treg studies, by better understanding the variable clinical association of Tregs across alternative tumor contexts. We show that increased expression of a 32-gene signature in Treg-enriched tumor samples (n=135) is able to distinguish a cohort of patients associated with chemosensitivity and overall survival This cohort is also enriched for CD8+ T cell abundance, as well as the antitumor M1 macrophage subtype. With a subsequent validation in a larger TCGA pool of Treg-enriched patients (n = 626), our results reveal a gene signature able to produce unsupervised clusters of Treg-enriched patients, with one cluster of patients uniquely representative of an immunogenic tumor microenvironment. Ultimately, these results support the proposed gene signature as a putative biomarker to identify certain Treg-enriched patients with immunogenic tumors that are more likely to be associated with features of favorable clinical outcome.

## Introduction

Studies of the tumor microenvironment have surfaced promising avenues of exploration to better understand the clinical relevance of T cell immunobiology. Regulatory T cells (Tregs) have keenly emerged in light of their ability to inhibit the adaptive immune response and provide a mechanism of immune escape for cancer cells within the tumor microenvironment across various cancer types [1–4]. However, the myriad of studies exploring the clinical relevance of intratumoral Treg abundance has produced controversial results to date [2], with some studies finding a poor prognosis associated with Treg infiltration [1, 5, 6], and others suggesting a favorable Treg-associated prognosis [6–12]. Accordingly, recent efforts to account for these polarized clinical results have undermined the notion that FOXP3+ Tregs invariably suppress tumor immunity [2, 13, 14]. Consequently, conducting additional unbiased analyses has been supported to better understand the clinical associations of Tregs across different microenvironmental contexts and tumor types[2, 14].

Using multiple gene markers together to accurately identify Tregs in cancer, such as FOXP3+BLIMP1[15] or FOXP3+CTLA4 [16], has demonstrated a more thorough evaluation of Tregs in recent years [13, 15]. Our current study also uses multiple markers to define Tregs enrichment rather than FOXP3 expression alone [17], but our primary interest was rather to examine the global microenvironmental context through which Tregs may be associated with differing clinical outcomes such as chemosensitivity and overall survival (OS). Moreover, we investigated whether these outcomes may be linked to CD8+ T cell abundance in pursuit of deriving a gene signature able to distinguish immunogenic Treg-enriched tumor samples. We hypothesized that a set of highly variable genes differentially expressed by Tregs (amongst 22 immune cell types) [18] would be able to produce distinct patient clusters from a pool of tumor samples that were all selected due to their enrichment for Tregs [17]. We implemented a pan-cancer approach to identify a favorable immunogenic signature based on immunological expression. In the current study, we present a 32-gene signature that is able to distinguish a ‘hot’ tumor phenotype associated with chemosensitivity, OS, and CD8+ T cell activation/abundance amongst a pool of 135 Treg-enriched patient tumor samples. We validated our 32-gene signature by confirming associations with OS, and CD8+ T cell activity/abundance in a larger pool of 626 patients. In addition, we overlapped an independently discovered set of genes suggested to be essential for CD8+ T cell function for immune checkpoint blockade therapy and assessed the concordance of the clusters produced by each gene set independently [19]. Together, we conducted a comprehensive expression analysis of a 32-gene signature and observed its association with favorable clinical features of tumor immunology.

## Results

### Unsupervised clustering using the 32 Treg DEGs

The experimental workflow is shown in Figure 1. Patient tumor samples analyzed in the current study were restricted to those that were (1) Treg-enriched (q < 0.05) and (2) possessed sufficient clinical drug response labels (n_Total_ = 135) (Figure 1) [17]. Using only the 32 highly variable genes differentially expressed by Tregs [18], k-means clustering was able to produce two distinct clusters of patient tumor samples (P = 0.0007, *x*^*2*^ = 11.58; Figure 2A). Cluster1 (n = 57) was enriched for chemosensitivity (70.17% of patients were sensitive to drugs prescribed), while cluster2 (n = 78) was enriched for chemoresistance (only 40.74% were clinically sensitive to drugs prescribed).

**Figure 1.**
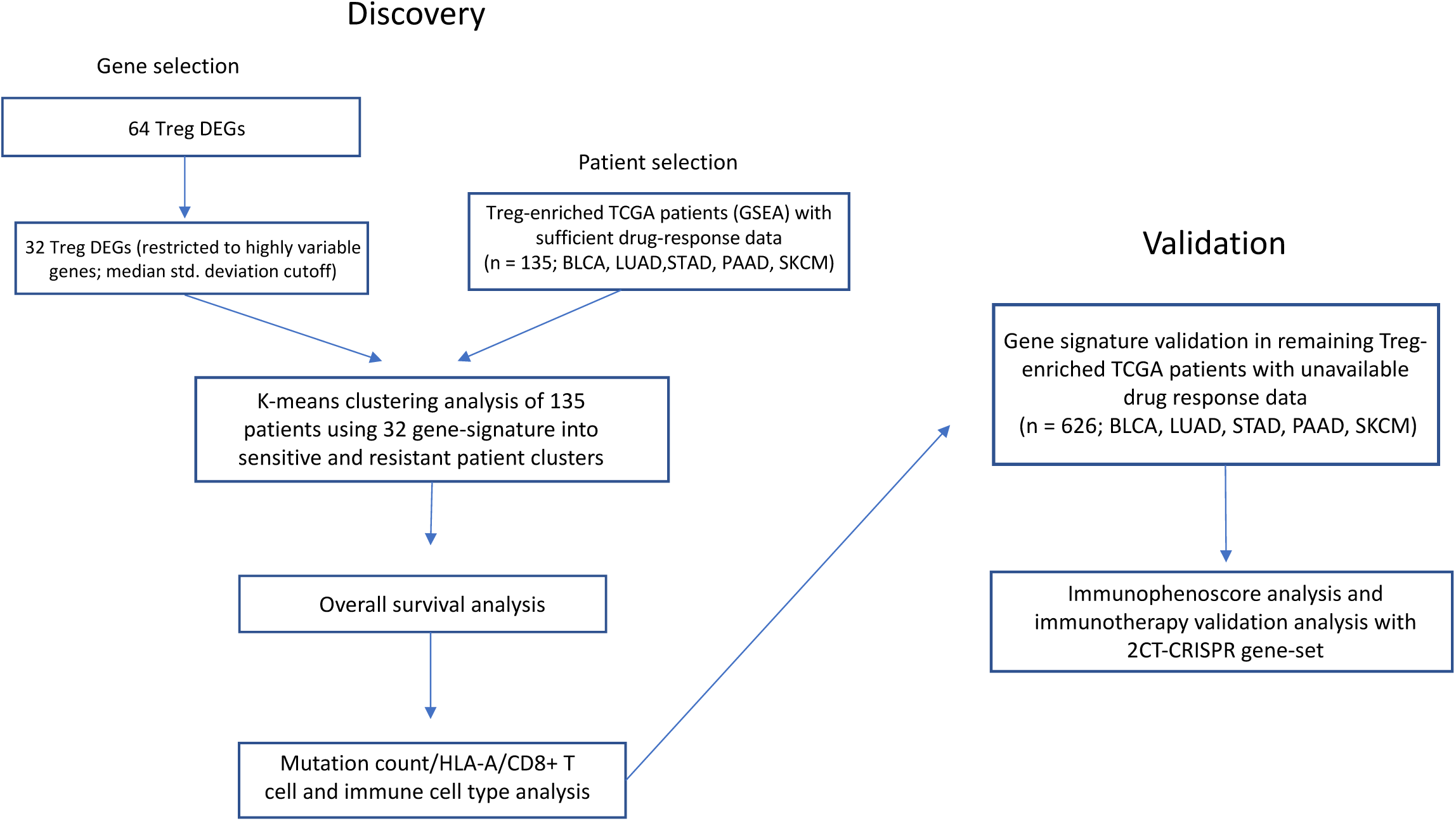
Experimental workflow of 32-gene signature expression analysis in discovery and validation cohorts of Treg-enriched cancer patients.

**Figure 2.**
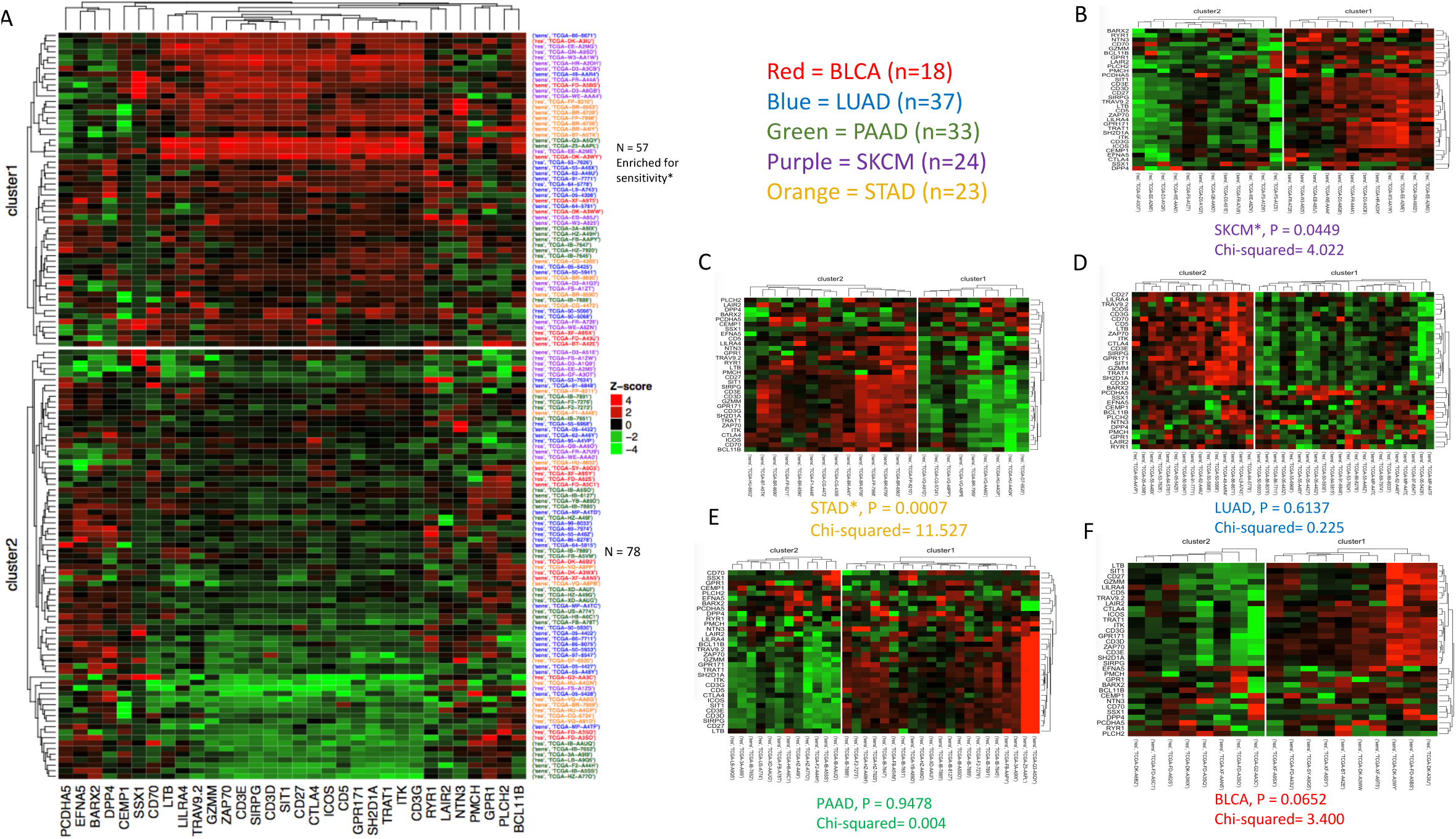
Unsupervised k-means clusters of Treg-enriched cancer patients from patient gene expressions of 32-gene signature in the discovery cohort (n = 135). (A) Clusters obtained from k-means clustering analysis (k=2), with cluster1 (n = 57, high/red) enriched for treatment sensitivity (P = 0.0007, *x*^2^ = 11.58) compared to cluster2 (n = 78, low/green), and neither cluster confounded with tumor anatomical location (P > 0.05; see Supp. Table 1-2). Row labels correspond to sensitivity marker (‘sens’ = Partial Response or Complete Response, ‘res’ = Stable Disease or Clinical Progressive Disease) and TCGA patient barcodes. Row label colors correspond to TCGA cancer type (18 BLCA [red], 37 LUAD [blue], 33 PAAD [green], 24 SKCM [purple], 23 STAD [orange]). (B) k-means clusters produced from 32-gene signature in SKCM (P = 0.0450, *x*^2^ = 4.022). (C) k-means clusters produced from 32-gene signature in STAD (P = 0.0007, *x*^2^ = 11.53). (D) k-means clusters produced from 32-gene signature in LUAD (P = 0.6137, *x*^2^ = 0.225). (E) k-means clusters produced from 32-gene signature in PAAD (P = 0.9478, *x*^2^ = 0.004). (F) k-means clusters produced from 32-gene signature in BLCA (P = 0.0652, *x*^2^ = 3.400).

**Table 1.** RNA-seq tumor sample deconvolution to immune cell type abundances (absolute value means) between clusters derived from 32-gene signature.

| Immune Cell Type | cluster1 mean abundances (n=57) | cluster2 mean abundances (n=78) | t-statistic | P-value |
| --- | --- | --- | --- | --- |
| T cells CD8 | 0.153530 | 0.073861 | 5.270214 | 0.000001 |
| Macrophages M0 | 0.108029 | 0.199764 | -4.258066 | 0.000038 |
| T cells CD4 memory activated | 0.021997 | 0.006819 | 4.209816 | 0.000069 |
| Neutrophils | 0.001244 | 0.010078 | -3.231105 | 0.001772 |
| Macrophages M1 | 0.072440 | 0.048957 | 2.745808 | 0.007010 |
| B cells naive | 0.103911 | 0.059839 | 2.372021 | 0.019814 |
| Mast cells activated | 0.001338 | 0.009605 | -2.288763 | 0.024394 |
| Dendritic cells activated | 0.014635 | 0.027883 | -2.245423 | 0.026440 |
| T cells regulatory (Tregs) | 0.020736 | 0.011808 | 2.249714 | 0.026875 |
| Macrophages M2 | 0.157614 | 0.192663 | -2.125468 | 0.035405 |
| Mast cells resting | 0.030808 | 0.044771 | -2.089000 | 0.038608 |
| NK cells activated | 0.006706 | 0.011616 | -1.460647 | 0.146820 |
| T cells follicular helper | 0.045859 | 0.036308 | 1.400419 | 0.163802 |
| T cells CD4 naive | 0.007653 | 0.000103 | 1.341638 | 0.185125 |
| Monocytes | 0.026926 | 0.038978 | -1.303338 | 0.195072 |
| Plasma cells | 0.038825 | 0.050363 | -1.160674 | 0.247865 |
| B cells memory | 0.038502 | 0.023567 | 1.160351 | 0.248782 |
| Dendritic cells resting | 0.005879 | 0.009324 | -1.080562 | 0.281837 |
| NK cells resting | 0.034977 | 0.033187 | 0.325962 | 0.745004 |
| T cells CD4 memory resting | 0.152862 | 0.149396 | 0.207072 | 0.836325 |
| Eosinophils | 0.000260 | 0.000223 | 0.159053 | 0.873866 |

Overall, patient cancer type was not associated with cluster (P > 0.05, *x*^2^ = 7.69; Supp. Table 1). Nevertheless, when the 32-gene signature was used to cluster patients of each cancer type individually, clusters enriched for sensitivity to treatment were observed in both SKCM and STAD (P = 0.0450, *x*^2^ = 4.022 and P = 0.0007, *x*^2^ = 11.53; Figure 2B-C). Proportional significance was not observed when identical clustering methods were applied independently to LUAD, PAAD, and BLCA (P = 0.614, P = 0.948, and P = 0.065, respectively; Figure 2D-F).

### Cluster1 patients are associated with favorable overall survival

To determine whether the cluster1 enrichment for drug-sensitivity was reflected in survival, overall survival (OS) survival analysis was performed (Figure 3). Cluster1 OS was observed to be 29.4 months, while cluster2 OS was observed shorter at 21.0 months, demonstrating an 8.4-month median survival difference. As hypothesized, cluster1 was significantly associated with favorable patient OS in comparison to cluster2 (P = 0.00084, HR = 0.40[0.24-0.70]; Figure 3A).

**Figure 3.**
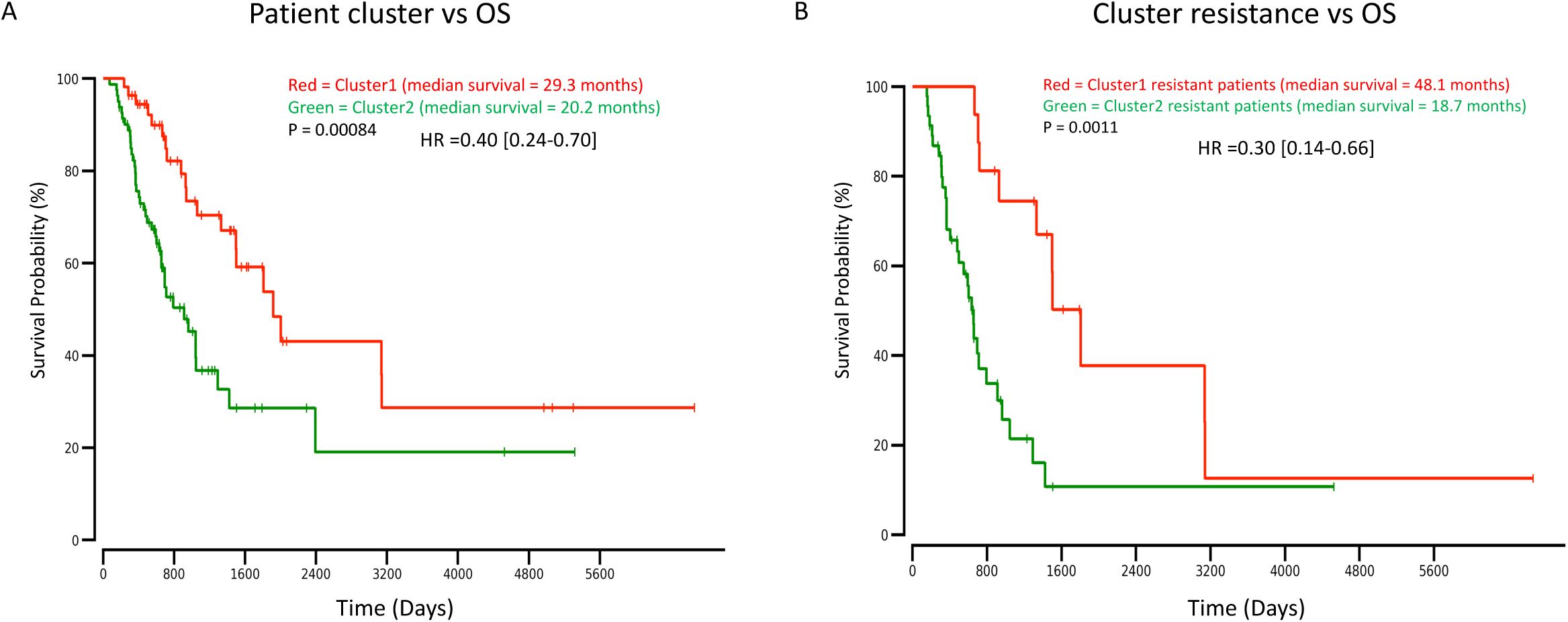
Overall survival plots/Cox proportional hazard regression analysis results. (A) OS analysis between cluster1 patients (n = 57, red) and cluster2 patients (n = 78, green), P = 0.00084, HR = 0.40[0.24-0.70]. (B) OS analysis between resistant patients from cluster1 (n = 17, red) and resistant patients from cluster2 (n = 45, green), P = 0.0011, HR = 0.30[0.14-0.66].

While cluster1 was enriched for patients who were sensitive to treatment, there was a handful of ‘resistant’ patients that were of further interest in this cluster (n = 17). To examine these patients further, we hypothesized that despite clinical labels for initial drug-response, patient OS would be significantly longer for an immunologically active ‘hotter’ tumor than it would be for a less immunologically active ‘colder’ tumor [19]. Therefore, we sought to determine whether ‘resistant’ patients in cluster1 differed in OS as compared to the ‘resistant’ patients of cluster2 (n = 45). After confirming that neither resistant patient cohort was enriched for a certain cancer type (P > 0.05; Supp. Table 2), we analyzed patient OS between each resistant cohort. Strikingly, the median survival duration of the resistant patients in cluster1 was more than twice that of the resistant patients in cluster2 (49.0 months vs 18.7 months, P = 0.0011, HR = 0.30[0.14-0.66]; Figure 3B).

### CD8+ T-cell activity/abundance is enriched in cluster1 patient tumor samples

To examine a putative explanation that may corroborate the observed clinical outcome parameters for the 32-gene signature (e.g., chemosensitivity and OS), we examined the distributions of mutation count per cluster and CD8+ T-cell biomarker expression, as mutation count and CD8/HLA-A expression have previously been observed in association with drug response and survival via neoantigen production and cytotoxic lymphocyte activation, respectively [20, 21]. To this end, and in line with this previous work, neither total mutation count nor non-synonymous mutation count was associated with cluster1 (P = 0.334 and P = 0.426; Supp. Fig 1A-B, respectively). However, CD8A/B, HLA-A, and PRF1 expressions were significantly elevated in patient tumor samples from cluster1 (P = 2.24e-19, P = 2.14e-18, P = 1.16e-4, and P =2.42e-10, respectively; Figure 4A-D).

**Figure 4.**
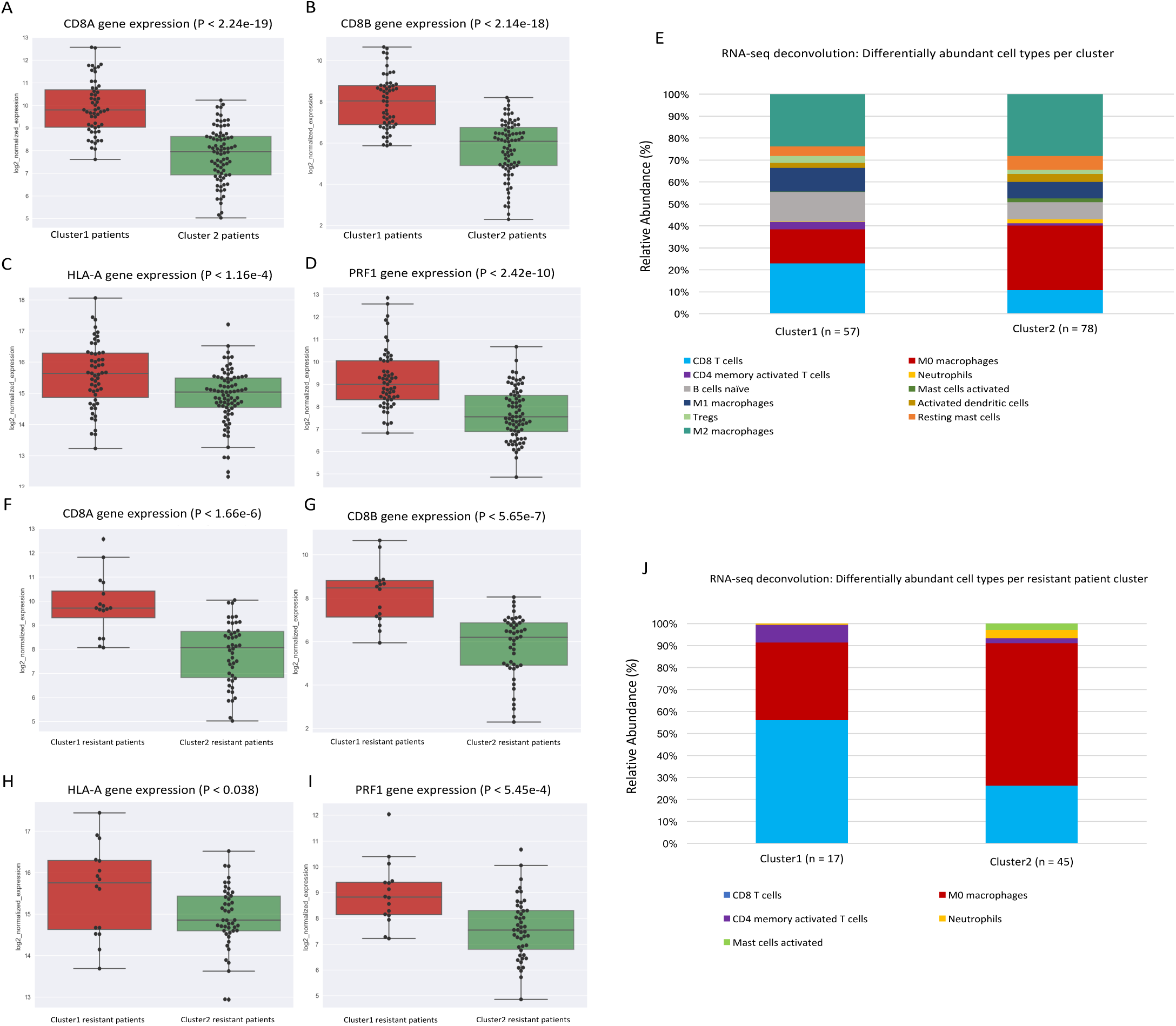
CD8+ T cell activity and immune cell abundance analysis between clusters. K-means clusters derived from 32-gene signature expression versus CD8+ T cell activity marker expression for (A) CD8A, (B) CD8B, (C) HLA-A, and (D) PRF1 (P = 2.24e-19, P = 2.14e-18, P = 1.16e-4, and P =2.42e-10, respectively; Figure 4A-D). (E) CIBERSORT RNA-seq deconvolution results for relative immune cell type abundances between patient tumor samples from k-means clusters (only differentially represented cell types (P < 0.05) were selected for presentation/visualization purposes; see Table 1 for absolute cell type abundances). Treatment resistant patients extracted from k-means clusters derived from 32-gene signature expression versus CD8+ T cell activity marker expression for (F) CD8A, (G) CD8B, (H) HLA-A, and (I) PRF1 (P = 1.66e-6, P =5.65e-7, P = 0.038, and P = 5.45e-4, respectively; Figure 4F-I). (J) CIBERSORT RNA-seq deconvolution results for relative immune cell type abundances between resistant patient tumor samples from k-means clusters (only differentially represented cell types (P < 0.05) were selected for presentation/visualization purposes; see Supp. Table 3 for absolute cell type abundances).

In order to suggest with further confidence at the level of immune cell type that CD8+ T-cells were more abundant in tumor samples from cluster1 patients, we applied a gene expression deconvolution framework to interpret the immunologically heterogeneous RNA-seq signals of each patient tumor sample [18]. In line with our observations of elevated CD8A/B gene expression in cluster1, CD8+ T-cells mean abundance was significantly higher in cluster1 patient tumor samples than that of cluster2 (15.3% vs 7.4% of total immune cells, P = 1.00e-7; Figure 4E and Table 1).

In parallel, continuing the analysis of cluster1 chemoresistant patients vs cluster2 chemoresistant patients, we examined mutation counts and expression of CD8A/B and HLA-A, as well as immune cell type abundances between these two resistant patient sub-cohorts. Similar to the previous broader cluster analysis, mutation count was not associated with chemoresistant patient cohort (P = 0.334 and P = 0.190; Supp. Fig 1C-D, respectively). Moreover, in line with our OS observations of these patients, CD8A/B, HLA-A, and PRF1 expressions were higher in resistant patient tumor samples from cluster1 than for those from cluster2 (P = 9.0e-6, P =2.0e-6, P = 0.038, and P=0.0008 respectively; Figure 4F-I). Furthermore, in resistant patients, CD8+ T-cells mean abundance was higher in cluster1 resistant patient tumor samples than that of cluster2 resistant patients (14.5% vs 7.6%, P = 0.013; Figure 4J, Supp. Table 3).

### Patient tumor sample DNA accessibility analysis

DNA accessibility refers to whether a certain region of DNA is accessible to regulatory molecules and proteins such as transcription factors, and recent work has revealed the importance of investigating the accessibility of regulatory regions during immune events such as CD8 T cell exhaustion, as accessible regions possess high potential to be expressed and or accessed by transcriptional machinery that may influence an immune response [22]. In line with this and due to the inherently high dimensional data of chromatin organization, deep learning models have been trained on DNase-seq [23] and recently extended to incorporate RNA-seq data to predict accessibility in a variety of cell types and tumor samples [24]. We therefore sought to explore whether cluster1 patients and cluster2 patients may differ in DNA site accessibility predictions across 86,057 promoter and promoter flank sites across the human genome. Interestingly, we observed sites within 14 genes that were predicted to be enriched for accessibility in cluster2 patients (acceptable P < 5.8e-7; Supplemental Figure 2A).

Of particular interest, we observed a site within TRAF3IP1 to be the most differentially accessible site of all 86,057 sites between clusters (P < 3.86e-9; Supplemental Figure 2C). TRAF3IP1 has been shown to negatively regulate the innate Type 1 IFN response [25], and indeed, TRAF3IP1 expression was elevated in cluster2 patients (P < 3.06e-4; Supplemental Figure 2B). Conversely, when investigating the pro-inflammatory immune marker IFNG [26, 27], we observed significant gene expression elevation in cluster1 patients (P < 1.78e-7; Supplemental Figure 2C). While of clear need for experimental validation, this suggests for the first time that TRAF3IP1 may be an unfavorable biomarker perhaps related to the anti-tumor immune response at least in Treg-enriched patient tumor samples.

### Gene signature analysis in validation cohort

In order to confirm that the application of this gene signature was not unique to the initial subset of patients selected, and that it extends to all Treg-enriched patients from the cohorts studied, we applied our 32-gene signature to the remaining Treg-enriched patients. For analytical consistency, these patients were those of the same cancer types (BLCA, LUAD, PAAD, SKCM, STAD) that were Treg-enriched (q < 0.05). However, these patients did not have available drug response data to assess treatment sensitivity.

Unsupervised clustering produced two clusters (n_cluster1_ = 332; n_cluster2_ = 294; Figure 5A). We observed that cluster1 patient tumor samples possessed significantly higher levels of CD8A, CD8B, HLA-A, and PRF1 (P < 4.19e-79, P < 1.59e-69, P < 2.96e-17, and P < 5.20e-58; Figure 5B-E, respectively). We also elaborated upon these CD8+ T cell activity markers by showing cluster1 patients possessed higher abundances of CD8+ T-cells than cluster2 patients (13.5% vs 6.90%; P = 2.15e-21; Figure 5F and Supp. Table 4).

**Figure 5.**
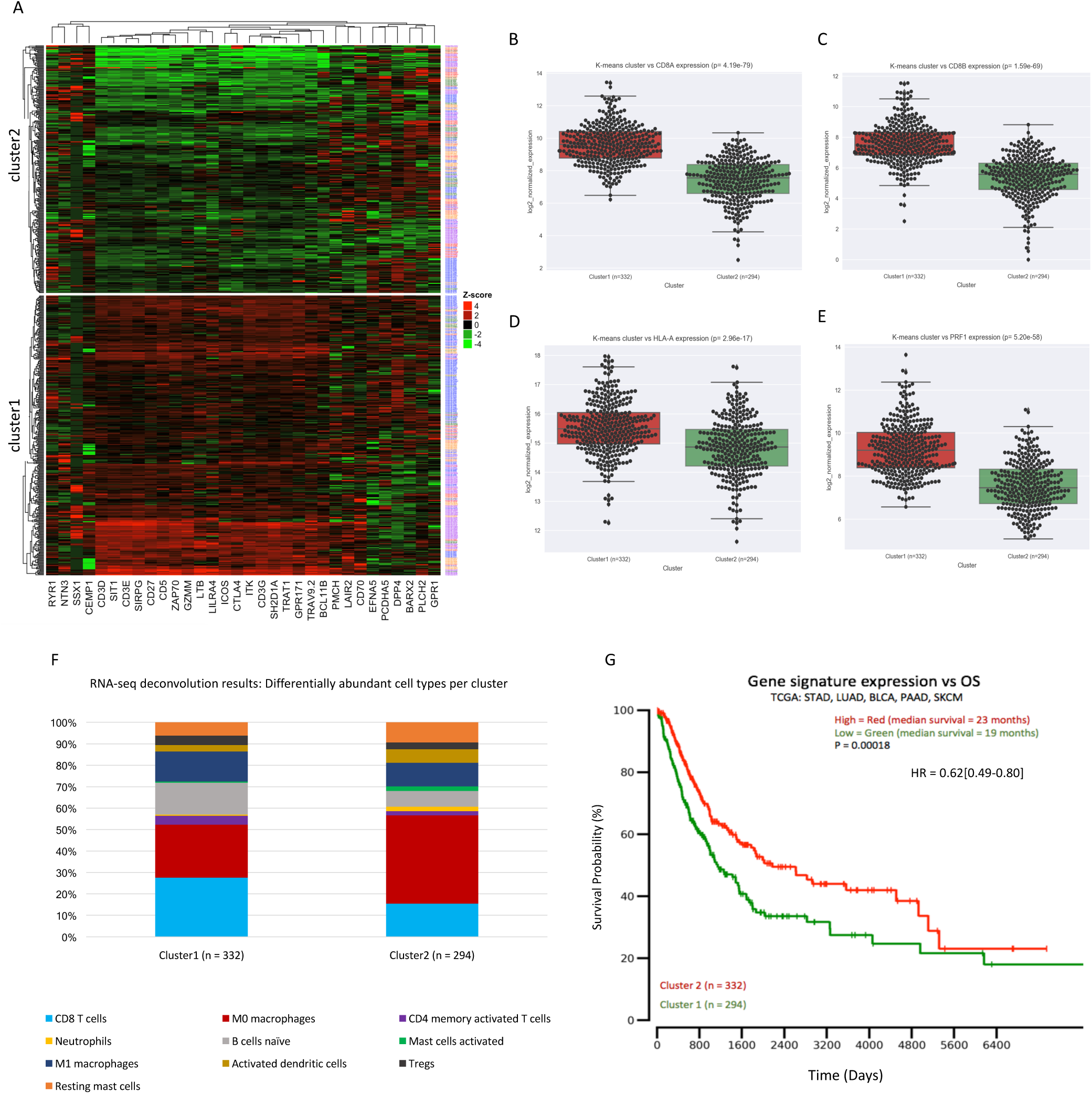
Gene signature analysis in TCGA validation dataset (n = 626). (A) 32-gene signature expression k-means clustering produces two clusters (n_cluster1_ = 332, n_cluster2_ = 294). Clusters derived from 32-gene signature expression in TCGA validation data versus CD8+ T cell activity marker expression for (B) CD8A, (C) CD8B, (D) HLA-A, and (E) PRF1 (P = 4.19e-79, P =1.59e-69, P = 2.96e-17, and P = 5.20e-58, respectively; Figure 4B-E). (F) CIBERSORT RNA-seq deconvolution results for relative immune cell type abundances between patient tumor samples from k-means clusters (only differentially represented cell types (P < 0.05) were selected for presentation/visualization purposes; see Supp. Table 4 for absolute cell type abundances). (G) Overall survival plot/Cox proportional hazard regression analysis results between cluster1 patients (red) and cluster2 patients (green), P = 0.00018, HR = 0.40[0.49-0.80].

Although clinical drug response data was not available for these 636 patients, we examined patient OS between the clusters produced from the 32-gene signature (Figure 5A). Indeed, higher expression of the gene signature was associated with OS in this larger dataset. The median OS duration for the higher expressing cluster1 patients was significantly longer than that of the cluster2 patients (P = 1.8e-4; Figure 5G), which was in line with the OS results previously discussed (Figure3A).

### Patient immunophenoscore and immunotherapy gene set concordance

Recent work has shown the utility of the Immunophenoscore (IPS) to predict response to immune checkpoint blockade in melanoma patient tumors based on higher pre-existing immunogenic potential [17]. We reasoned that if cluster1 was indeed representative of a more immunogenic phenotype, then cluster1 tumor samples should too display an elevated IPS, which has been clinically validated for immunotherapeutic response [17]. Analysis of patient IPS between cluster1 and cluster2 revealed significant elevation in the general IPS of cluster1 (P = 0.019). In addition, the IPS-PD1, IPS-CTLA4, and IPS-PD1+CTLA4 scores were significantly elevated in cluster1 tumor samples (P = 3.68e-10, P = 6.00e-6, and 5.52e-12, respectively; Figure 6A). These three scores were designed to be assessed for patients with the potential to be administered checkpoint blockade therapy (e.g., Nivolumab, Ipilimumab), who possessed elevated expression of PD-1 and or CTLA-4 [17]. Indeed, cluster1 patients possessed significantly higher PD-1 and CLTA-4 gene expression than did cluster2 patients (P < 8.34e-15 and P < 1.83e-13; Figure 6B-C, respectively).

**Figure 6.**
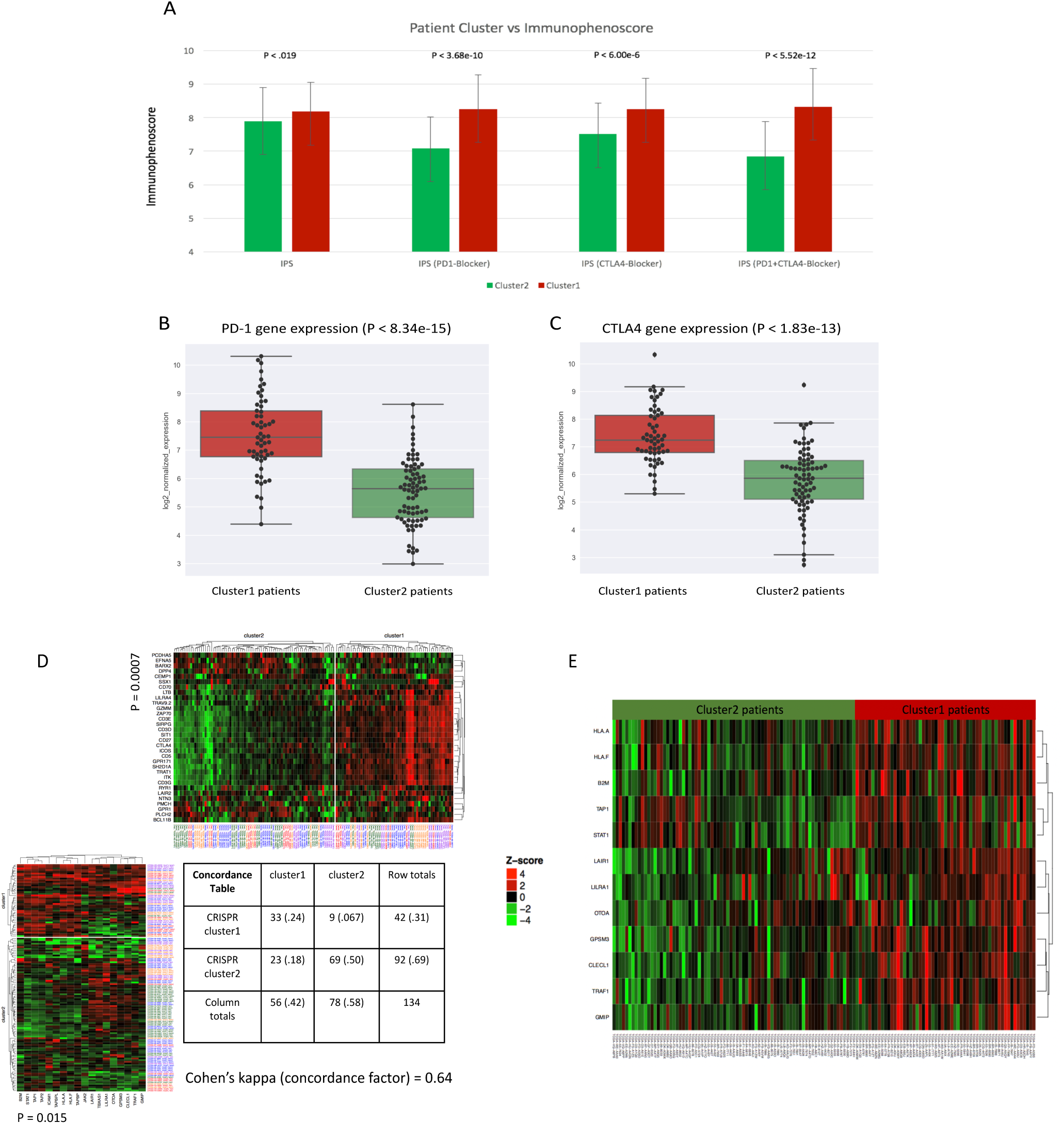
Patient immunophenoscore and immunotherapy gene set analysis. (A) mean Immunophenoscores represented per cluster derived as described in methods (bar whiskers represent standard deviations). Normalized gene expression plots between cluster1 and cluster2 patients for (B) PD-1 (P < 8.34e-15) and (C) CTLA-4 (P < 1.83e-3). (D) K-means clusters produced by each the immunotherapy gene set (19) and our gene set, with both gene sets showing enrichment for chemosensitive tumor samples (P = 0.015, and P = 0.0007, respectively). Concordance table showing similar unsupervised clusters are produced with ‘good’ concordance (Cohen’s kappa = 0.64). (E) cluster1 patients possessed significantly elevated expression for 12 out of 18 genes identified by the 2CT-CRISPR assay system (acceptable P < 0.003).

In order to lend further support to our 32-gene signature, we measured the significance of and concordance with another gene set recently suggested to be essential for effector CD8+ T cell activity for immunotherapy [19]. This gene set was identified by a 2CT-CRISPR assay as candidate genes essential for effector CD8+ T cell function and were correlated with cytolytic activity across almost all TCGA cancer types. When we applied k-means clustering, this immunotherapy gene set achieved significance in terms of enrichment for chemosensitive tumor samples (P = 0.015; Figure 6D), although this was less significant than that of our 32-gene signature (P = 0.0007; Figure 6D). More importantly, however, the concordance or agreement between patient tumor sample clustering by each gene set independently was ‘good’ (Cohen’s kappa = 0.64; Figure 6D)[28, 29], suggesting that our 32-gene signature was able to produce patient tumor sample clusters in concert with those experimentally derived from an immunotherapy gene set important for CD8+ T cell function [19].

Moreover, and in line with our k-means clustering results, we observed that cluster1 patients possessed significantly elevated expression for 12 out of 18 genes identified by the 2CT-CRISPR assay system (acceptable P < 0.003; Figure 6E). Interestingly, pathway network activation revealed cluster1 patients to possess enriched immune pathway activation via elevated T-Cell Receptor and Co-stimulatory signaling, TCR Signaling pathway, and the Inflammatory Response Pathway (P < 0.048, P < 0.048, and P < 0.043, respectively; Supp. Figure 3A), and these pathways were not enriched in cluster2 patients (Supp. Figure 3B). Together these results lend support to our gene signature as a clinically relevant gene expression signature of a ‘hot’ immunogenic tumor in Treg-enriched patients who may be more likely to respond to checkpoint blockade therapies targeting PD-1 and or CTLA-4.

## Discussion

Our study examined the clinical potential of a 32-gene expression signature to classify patients and their association with chemosensitivity, OS, and CD8+ T cell activation/abundance. The genes chosen to comprise this signature were differentially expressed by Tregs [18] and were highly variable across Treg-enriched patient tumor samples. We observed that the variation of these genes to corresponded to the ‘hot’ vs ‘cold’ tumor paradigm, in which a ‘hot’ tumor with a higher level of tumor infiltrating lymphocytes (TILs) is associated with favorable clinical outcomes[12, 30–34]. From our results, it is worth noting that in addition to an immunogenic tumor microenvironment, the tumor gene signature we investigated may be associated with a previously observed molecular phenotype of Tregs associated with diminished suppressive function [35].

Despite the proportional significance achieved by the number of chemosensitive patients between cluster1 and cluster2, there remained 17 chemoresistant patients in cluster1. To interrogate these patients further, we conducted a series of parallel analyses between these 17 patients and the 45 resistant patients of cluster2. Strikingly, we observed that although labeled as chemoresistant, the cluster1 resistant patients likely possessed ‘hotter’, more immunogenic tumors, and thus survived longer in terms of overall survival. Specifically, we observed favorable tumor infiltrating lymphocyte expression in these 17 tumor samples from our deconvolution analysis (e.g., more CD8+ T cells, less M0 macrophages) and augmented CD8+ T cell activation expression (e.g., CD8A/B, PRF1, HLA-A). Together, these clinical parameters agreed with associated patient OS findings. Our results suggest that at least in a small proportion of Treg-enriched patients, the ‘heat’ of a tumor classified by our 32-gene signature may complement the clinical parameter of chemosensitivity, and that the chemoresistant patients we studied may have been promising candidates for immunotherapy. It is also interesting to note that while checkpoint marker expression (e.g., PD-1 and CTLA-4) was highly elevated in cluster1 patients, PD-L1 expression was only marginally elevated (P < 3.6e-2, data not shown). This difference in significance between PD-1 and PD-L1 expression can perhaps be explained by the initial Treg-enrichment filtering.

When differential DNA accessibility was examined between cluster1 patients and cluster2 patients, a promoter flank site within the TRAF3IP1 gene was predicted to be accessible in almost all cluster2 patients and inaccessible in almost all cluster1 patients. TRAF3IP1 is a protein that interacts with TRAF3 and is has been observed to inhibit the innate type I IFN response [25].

TRAF3IP1 was also higher expressed in cluster2 patients who had lower expression of IFNG and anti-tumor immune activity, suggesting a potential role for TRAF3IP1 regulation in inhibiting the anti-tumor immune response. This, however, would require a thorough experimental validation to hold preliminary clinical relevance. Additionally, it is worth noting that the greatest visible difference in accessibility was specifically between melanoma patients of cluster1 and melanoma patients of cluster2, which is in line with previous work that demonstrated accessibility playing a role in CD8+ T cell immunoreactivity within a melanoma model [22].Furthermore, recent work in the B16 melanoma model has argued for the rational of coupling HDAC inhibitors to checkpoint blockade therapies to enhance immunotherapy efficacy [36]. This may further propose a clinical relevance for DNA accessibility to be explored in future studies.

While a literature search to biologically explain each gene of the 32-gene signature was not part of the current study, one gene of that particularly struck interest was Lymphotoxin Beta (LTB), which was elevated in cluster1 patient tumor samples. Interestingly, LTB has been shown to specifically stimulate Tregs to migrate from the tissue to the lymph nodes via afferent lymphatics [37]. This therefore may at least partly attribute the positive clinical associations of cluster1 patients to elevated LTB expression, which may be directing immunosuppressive Tregs away from tumor tissue sites, thus unleashing the antitumor response within cluster1 tumor samples. Clearly, this speculation would require further *in vivo* tumor studies to be confirmed.

In order to obtain sufficient sample sizes, we used a pan-cancer approach to interrogate differences between the clusters produced by the unsupervised k-means method. We confirmed that neither cluster was enriched for any single cancer type, which may have otherwise surfaced the tumor tissue type as a confounding variable. We additionally pursued an extensive analysis to deconvolute the heterogeneity of the tumor, showing that CD8+ T cell activity was likely enriched in cluser1 patient tumor samples and that chemosensitivity was seemingly linked to the immunocompetence of the ‘hot’ tumors [38]. To this end, we observed CD8+ T cell tumor abundance to be associated with favorable patient prognosis [38–42]. Of note, the pro-inflammatory IFNγ, which is a marker of CD8+ T cell mediated tumor regression and TIL abundance[43], was also significantly elevated in cluster1 samples. Although all patient tumor samples of cluster1 were also enriched for Treg expression, cluster1 possessed a higher mean CD8+ T cell to Treg ratio than cluster2 (data not shown). This result, which is based on the expression of our 32-gene signature, is also in line with previous clinical findings pertaining to the CD8+ T cell to Treg ratio in the tumor microenvironment [38, 44]. Furthermore, we observed that FOXP3 expression was significantly elevated in the clinically favorable cluster of patient tumor samples (cluster1, P < 5.41e-14; data not shown), which is also in accord with previous FOXP3 related studies [45–49].

While we specifically interrogated CD8+ T cell abundance/activation markers, we also observed differential activation of tumor associated macrophages (TAMs), with cluster1 patients enriched for M1 macrophage abundance. The M1 subtype, which is associated with higher levels of IL-1, TNFa, IL-12, and CXCL12, is associated with the inflammatory, anti-tumor response [50–52]. Our observation was therefore in line with previous work, suggesting a complementary interaction between M1 macrophages and CD8+ T cells to ameliorate pro-tumorigenic activity within cluster1 patient tumor samples. Cluster2 accordingly possessed a higher abundance of M0 macrophages, potentially suggesting a larger undifferentiated premature pool of monocytes in these patients as opposed to those of cluster1. Our pathway network activation analysis from a more holistic perspective also revealed a shift towards activated TCR signaling and the inflammatory response in cluster1 patients but not in cluster2 patients. Together, we observed that our 32-gene feature set was able to distinguish a drug-sensitive cohort (cluster1) with higher antitumor immune activity perhaps in part via the previously proposed interplay between TAMs and CD8+ T cells [53]. It is also interesting to note that the cluster2 patient pool (which was enriched for chemoresistance) uniquely possessed upregulated pathway activation for DNA damage response and nucleotide metabolism (P < 0.037 and P < 0.026, respectively). In agreement with previous work, these molecular processes have been observed to play a role in chemoresistance and survival likely by ameliorating the intended damage of chemotherapeutic compounds [54–58].

There are several limitations to our study. First, our 32-gene signature is specific is to Treg-enriched patient tumor samples as determined by the previously described GSEA methods [17]. Second, the small number of Treg-enriched patients with sufficient drug response data was limited (n=135) and thus prevented us from establishing a robust aggregate cutoff to examine in a held-out validation dataset. To partially address this, however, we used an unsupervised clustering method which rendered our clusters independent of clinical variables (e.g., tumor stage, lymph node status, age), and we showed that our clusters are in good concordance with clusters derived from an independently proposed set of genes important for antitumor CD8+ T cell function. Third, as both a strength and a limitation, our chemosensitivity analysis was not specific to a certain drug, but rather for response to treatment for BLCA, LUAD, PAAD, SKCM, and STAD patients. Although this makes follow-up studies pertaining to a specific chemotherapeutic compound for a certain clinical indication less obvious, it allowed us to corroborate the paradigm of a ‘hot’ tumor across five tumor types and propose an immunogenic gene expression signature independent of clinical variables.

In conclusion, our study proposes a clinically robust 32-gene signature able to distinguish patients with a favorable phenotype in part through CD8+ T cell activity and abundance. Through a pan-cancer analysis of Treg-enriched patient tumor samples, we show two opposing contexts in which Tregs may be associated with alternative clinical features. This study marks the first pan-cancer patient study of its kind to interrogate tumor transcriptomic data to help reconcile the growing controversy of Tregs and their clinical impact. We believe the proposed gene signature may also serve as a promising starting point to extrapolate disease-specific gene signatures that could further sculpt the landscape of tumor immunobiology.

## Methods

### Patient Selection and drug response data

We used curated records of drug treatments and outcomes generated from TCGA clinical data [59] to treatment sensitivity labels. To analyze patient tumors overtly sensitive to treatment, patients with ‘Complete Response’ or ‘Partial Response’ were assigned to a ‘sens’ label, while patients with ‘Stable Disease’ or ‘Clinical Progressive disease were assigned to a ‘res’ label (Figure 2). Sensitivity to treatment served as one parameter to assess clustering output significance (further discussed below).

RNAseq data (TOIL RSEM norm_count) was downloaded and used as input features for unsupervised clustering was from the TCGA Pan-Cancer cohort from xenabrowser.net (dataset ID: tcga_RSEM_Hugo_norm_count, unit: log2(norm_count + 1), hub: GA4GH (TOIL) hub). This dataset contained 10,535 tumor samples, but only those with available drug response data enriched for Tregs were examined. We executed the filtering for Treg-enriched tumor samples via The Cancer Immunome Database (tcia.at) using gene set enrichment analysis (GSEA) of a non-overlapping, pan-cancer derived set of genes representative for Treg enrichment (FOXP3, CCL3L1, CD72, CLEC5A, ITGA4, L1CAM, LIPA, LRP1, LRRC42, MARCO, MMP12, MNDA, MRC1, MS4A6A, PELO, PLEK, PRSS23, PTGIR, ST8SIA4, STAB1). This method has been previously described and clinically validated in further detail [17]. For a TCGA cohort to be included in our analysis, it was required to comprise samples that were (1) enriched for Tregs (q < 0.05) [17], and (2) have sufficient clinical drug response data (at least 15 samples with available drug response labels in each cohort). Coalescing and applying these two filters yielded 135 total patients for analysis across 5 TCGA cohorts (18 BLCA, 37 LUAD, 33 PAAD, 24 SKCM, 23 STAD). This allowed a sufficient population for a pan-cancer unsupervised gene-signature expression analysis (n=135).

### Gene signature selection and patient clustering

K-means clustering is a popular vector quantization method that we used to classify the 135 patient tumor samples (k=2) via the ComplexHeatmap package in R using the Treg differentially expressed genes (DEGs) previously produced [18]. However, instead of using all 64 Treg DEGs available, we used only the highly variable 32 genes as our feature gene set (outlier(s) removed and median standard deviation cutoff for 64 genes expressions within the 135 patient tumor samples). This 32-gene set served as the feature set we used for k-means clustering (Figure 2) (for clustering visualization, the expression of each gene was scaled to the z-score relative to expression of that gene across the 135 tumor samples). As assumed by the unsupervised classification process, drug response labels were not included as input features, and nor were cancer types. Together, the input dataset consisted of a table with 135 rows (TCGA patient IDs) and 32 columns (32 highly-variable Treg DEGs) of z-score scaled expression values. These procedures were replicated for the 626 patients of the validation data set.

Proportional cluster significance for patient drug response sensitivity enrichment was determined using the “N-1” Chi-squared test (DF =1, P < 0.05 was considered significant) [60, 61]. To eliminate the possibility of cancer-type serving as a confounding variable, and thus ensuring that cohorts analyzed were not unevenly representative of a certain cancer type, a Chi-squared tests of independence in a contingency table was performed using the scipy.stats module in Python (P < 0.05 was considered significant).

### Survival analysis

Overall survival (OS) data for the 135 patients was downloaded from OncoLnc (oncolnc.org) [62]. Survival datasets for the BLCA, LUAD, PAAD, SKCM, and STAD cohorts were parsed and Kaplan-Meier plots were generated in Python. Log-rank tests were conducted using the Lifelines implementation in Python (lifelines 0.11.1) (1 month was considered 30 days; P < 0.05 was considered significant).

### Genomic/transcriptomic analysis

Patient genomic mutations and transcriptomic expression values were accessed using the fbget Python API and parsed in Python (https://confluence.broadinstitute.org/display/GDAC/fbget). Mutations were retrieved via whole-exome-sequencing MAF data, and gene expression values were retrieved as log2-normalized values via RSEM. The Mann-Whitney test was performed to compare mutation counts between patient clusters/cohorts. Two-tailed t-tests were used to determine significantly differential expression of CD8A, CD8B, and HLA-A between cohorts [21]. For both mutation count and differential gene expression, P < 0.05 was considered significant. Mutation histograms and expression swarm-boxplots were generated in Python using the Matplotlib and Seaborn libraries.

### Immune-cell type abundance

To characterize immune cell composition within the 135 patient tumor samples, we used CIBERSORT [18]. We used this method to present cell-type abundance via RNA transcript expression from a recommended gene expression profile of genes for 22 immune cell types. The recommended model parameters were used for this prediction task (e.g., LM22 signature gene file, disabled quantile normalization for RNA-seq data), and 1,000 permutations were conducted [18]. In short, this strategy applies nu-support vector regression (*v*-SVR) to discover a hyperplane that separates classes of interest. CIBERSORT has been shown to outperform other approaches and has been previously described in further detail [18].

For upregulated pathway network analysis, we used the integrated software workflow AltAnalyze, which is tool is used to analyze expression data in the context of interaction networks and pathways [63, 64]. The input data consisted of whole-transcriptome data from each patient (with 3,325 tissue-specific genes from TissGDB filtered out [65] due to the pan-cancer methodology) obtained from TOIL RSEM data [66] and encoded as log2(TPM + 1) with no other normalizations. Enriched/upregulated pathways (from WikiPathways [67]) were those with at least a 2.0-fold-change surfaced from the integrated GO-Elite software that retained significance (P < 0.05) after Benjamini-Hochberg Fisher’s exact P-value adjustment.

### DNA accessibility analysis

DNA accessibility predictions across whole genomes for TCGA patients of interest were generated using a convolutional neural network model that has recently been shown to make accurate predictions at promoter and promoter flank sites, even for novel biotypes not seen in training [24]. The model makes binary accessibility predictions for 600-base-pair DNA sequences centered at potentially accessible sites, as identified through agglomerative clustering. Model input includes the DNA sequence of a specific site augmented with an input vector of select gene expression levels estimated from RNA-seq. This additional RNA-seq vector is what makes it possible for the neural network to learn to modulate accessibility predictions appropriately according to the cellular context and make predictions for unseen cell types.

Gene expression levels for patient samples were obtained from TOIL RSEM data [66] and encoded as log2(TPM + 1) with no other normalizations. Since whole genome sequencing was not available for all patients of interest, all predictions were made using the reference genome hg19/GRCh37. It is therefore possible that some differences between patients were missed due to this lack of mutation information. Empirically it was found that including mutation information affected the classification outcome at 5.5% of promoter and promoter flank sites.

The model was trained to make predictions at 1.71 million genomic sites, with a training set consisting of 338.7 million training examples, spanning 66 unique cell and tissue types from ENCODE [24]. Of the 1.71 million training sites, the TCGA predictions were restricted to 108970 sites that overlapped with promoter and promoter flank annotations. A classification threshold was selected such that the neural net achieved 80% precision on those sites when making predictions for novel biotypes, at which it demonstrated a recall of 65.3% and false positive rate of 10%. For analysis only the subset of those promoter and promoter flank sites that overlapped with protein coding genes (as annotated by GENCODE v19 and extended by 1k base pairs front the TSS) were considered, reducing the number of regions to 86,057.

### Immunophenoscore and immunotherapy gene analysis

As previously described [17], a patient’s immunophenoscore (IPS) can be derived in an unbiased manner using machine learning by considering the four major categories of genes that determine immunogenicity (effector cells, immunosuppressive cells, MHC molecules, and immunomodulators) by the gene expression of the cell types these comprise (e.g., activated CD4+ T cells, activated CD8+ T cells, effector memory CD4+ T cells, Tregs, MDSCs). The IPS is calculated on a 0-10 scale based on representative cell type gene expression z-scores, where higher scores are associated with increased immunogenicity. This is because the IPS is positively weighted for stimulatory factors (e.g., CD8+ T cell gene expression) and negatively weighted for inhibitory factors (e.g., MDSC gene expression). Finally, the IPS is calculated based on a 0-10 scale relative to the sum of the weighted averaged z-scores. A z-score of 3 or more translates to an IPS of 10, while a z-scores 0 or less translates to an IPS of 0, demonstrating a higher IPS is representative of a more immunogenic tumor [17]. This method has been described in further detail with the immunogenic determinant categories, as well as corresponding cell types and gene sets, which can be found at tcia.at [17].

We retrieved patient IPSs from The Cancer Immunome Atlas framework. Relative bar plots were generated for visualization and error bars reflect standard deviations. Two-tailed t-tests were used to determine significantly differential IPS values (P < 0.05 was considered significant).

The 19 genes used to reinforce our 32-gene signature derived clusters were taken from a previous study that used a 2CT-CRISPR assay system to identify 554 candidate genes essential for immunotherapy[19]. Via hierarchical clustering, the gene set we examined were those of the 554 genes that identified to be correlated with cytolytic activity across most TCGA cancer types [19]. One patient sample and one gene were lost while conducting this analysis due to data availability (e.g., 18 genes and 134 patients were examined). To measure concordance, we calculated Cohen’s kappa via a concordance table of clusters predicted from k-means with our 32-gene set and the Patel 18-gene set [29]. A Cohen’s kappa value of 0.61-0.80 constituted a ‘good’ quality of agreement, as previously described [28, 29].

The heatmap representative of the relative expressions of the 12/18 genes significantly elevated in cluster1 patients (acceptable P < 0.003) was produced with the ComplexHeatmap package in R. Clusters represented are those derived from our initial 32-gene k-means clustering. We used this gene set as validation to show that the 32-gene signature used for clustering was able to produce a cluster with high expression of the same genes recently identified as likely essential for immunotherapy in an independent setting.

